# Paternal genome elimination creates contrasting evolutionary trajectories in male and female citrus mealybugs

**DOI:** 10.1101/2024.07.28.604693

**Authors:** Andrew J. Mongue, Tamsin Woodman, Hollie Marshall, Arkadiy Garber, José C. Franco, John P. McCutcheon, Laura Ross

## Abstract

Most studies of sex-biased genes explore their evolution in familiar chromosomal sex determination systems, leaving the evolution of sex differences under alternative reproductive systems unknown. Here we explore the system of paternal genome elimination employed by mealybugs (Hempitera: Pseudococcidae) which have no sex chromosomes. Instead, all chromosomes are autosomal and inherited in two copies, but sex is determined by the ploidy of expression. Females express both parental alleles, but males reliably silence their paternally inherited chromosomes, creating genome-wide haploid expression in males and diploid expression in females. Additionally, sons do not express alleles directly inherited from their fathers, potentially disrupting the evolution of male-benefitting traits. To understand how these dynamics impact molecular evolution, we generated sex-specific RNAseq, a new gene annotation, and whole-genome population sequencing of the citrus mealybug, *Planococcus citri*. We found that genes expressed primarily in females hold more variation and evolve more quickly than those expressed in males or both sexes. Conversely, more adaptation occurs in genes expressed mainly in males than those expressed in females. Put together, paternal genome elimination appears to slow change on the male side but, by increasing selective scrutiny, increase the amount of adaptation in these genes. These results expand our understanding of evolution in a non-mendelian genetic system and the data we generated should prove useful for future research on this pest insect.

## Introduction

Foundational to the study of evolutionary genetics is understanding the forces that drive sequence change in organisms. These can be described in the simplest terms as either random change through genetic drift, or non-random change via selection; however, this simple binary belies the fact that molecular evolution is the amalgamation of numerous biological factors, both random and selective, acting on the organism expressing genotypes as phenotypes. Thus, a more nuanced statement of the goal of evolutionary genetics is to unpack the numerous, often conflicting pressures on gene evolution to understand which factors are the key drivers of change, and under what conditions.

A prime example of this framework is the study of sex-biased gene evolution (Meisel 2011). Males and females necessarily have phenotypic differences, so expression of male phenotypes in a female (and vice versa) is often costly; thus, these genes face an inherent conflict: being beneficial in one sex but deleterious in the other (explored extensively in Arnqvist and Rowe 2005). Constraining this conflict is the fact that, for the most part, males and females of the same species share the same genome.

Consequently, males and females each hold genes for both sexes within their genome, meaning a male-benefitting gene will be carried in the genome of a female at some point (and vice versa). The long term evolutionary resolution to this conflict is conditional, or sex-biased, expression of genes; male-benefitting genes become male-biased in expression and female-benefitting genes become female-biased (Wright et al. 2019). Although this solution better aligns gene expression with the sex that most benefits, conditional expression creates new countervailing forces and evolutionary genetic contradictions.

On the one hand, limiting expression to one sex decreases the effective population size in which these genes are exposed to selection. Under the framework of the nearly neutral theory of molecular evolution, this reduction should make selection less efficient and increase fixation of nonadaptive alleles through drift for sex-biased genes compared to those expressed in both sexes (Ohta 1992; Baines et al. 2008; Dapper and Wade 2016). On the other hand, many of these biased genes encode reproductive traits and are often observed or predicted to evolve more quickly and adaptively than other genes thanks to their roles in species boundary formation and sexual selection (Civetta and Singh 1995; Swanson and Vacquier 2002; Wright et al. 2015). Thus, understanding which factors are most important to the evolution of sex-biased genes is challenging to say the least.

In practice, the study of sex-biased gene evolution is usually associated with the study of sex chromosomes (Ellegren 2011; Albritton et al. 2014; Sackton et al. 2014) in part for the practical reason that these chromosomes tend to be enriched for sex-biased genes compared to the autosomes (Allen et al. 2013; Jaquiéry et al. 2013; Mongue and Walters 2017). And indeed, there is a wealth of research exploring the role of adaptation and drift in the evolution of sex-biased genes on the sex chromosomes (Mank et al. 2009; Meisel and Connallon 2013; Dean et al. 2015; Rousselle et al. 2016; Whittle et al. 2020; Mongue et al. 2022; Mongue and Baird 2024). Although patterns like increased rates of molecular evolution of sex-biased genes have emerged in the study of sex chromosomes, they also come with a number of confounding biological factors that set them apart from the autosomes including smaller population sizes in any given species (Vicoso and Charlesworth 2009; Mank et al. 2010), sex-biased recombination rates in some taxa (Turner and Sheppard 1975; John et al. 2016; Danzmann et al. 2019), and variable gene regulation compared to the autosomes (Disteche 2012; Gu and Walters 2017). More systematic study of sex chromosomes in a wider array of taxa will doubtless help understand the interplay of these facets, but to better understand what factors most impact sex-biased genes, it would be valuable to explore their evolution under a variety of genomic architectures. In particular, we would like to introduce a new model, the citrus mealybug, to explore the evolution of sex-biased genes in the absence of sex chromosomes.

### The unique genetics of the mealybug model system

The group containing the citrus mealybug, the scale insects (Hemiptera: Coccoidea), likely possessed X sex chromosomes ancestrally, as other Hemiptera and early diverging scale insects still employ this sex determination mechanism (Nur 1980; Gavrilov 2007; Blackmon et al. 2017). But many scale insects, including the citrus mealybug, *Planococcus citri*, employ an unusual alternative to sex chromosomes known as the paternal genome elimination (PGE) system of sex determination (Nur 1980; Tree of Sex Consortium 2014; Blackmon et al. 2017; Ross et al. 2022). In this system, males and females share the entirety of their genome, i.e., all chromosomes are autosomal and found in both sexes, but differ in the number of copies expressed in and passed on to offspring (Figure 1a). The initial sex difference is that embryos that develop as male transcriptionally silence their paternally inherited alleles (Nelson Rees 1962; de la Filia et al. 2021) and fail to pass these silenced alleles onto the next generation; females, in contrast, express and transmit both maternally and paternally inherited alleles (Schrader 1921; Hughes-Schrader 1948; Nur 1966; Herrick and Seger 1999).

**Figure 1.**
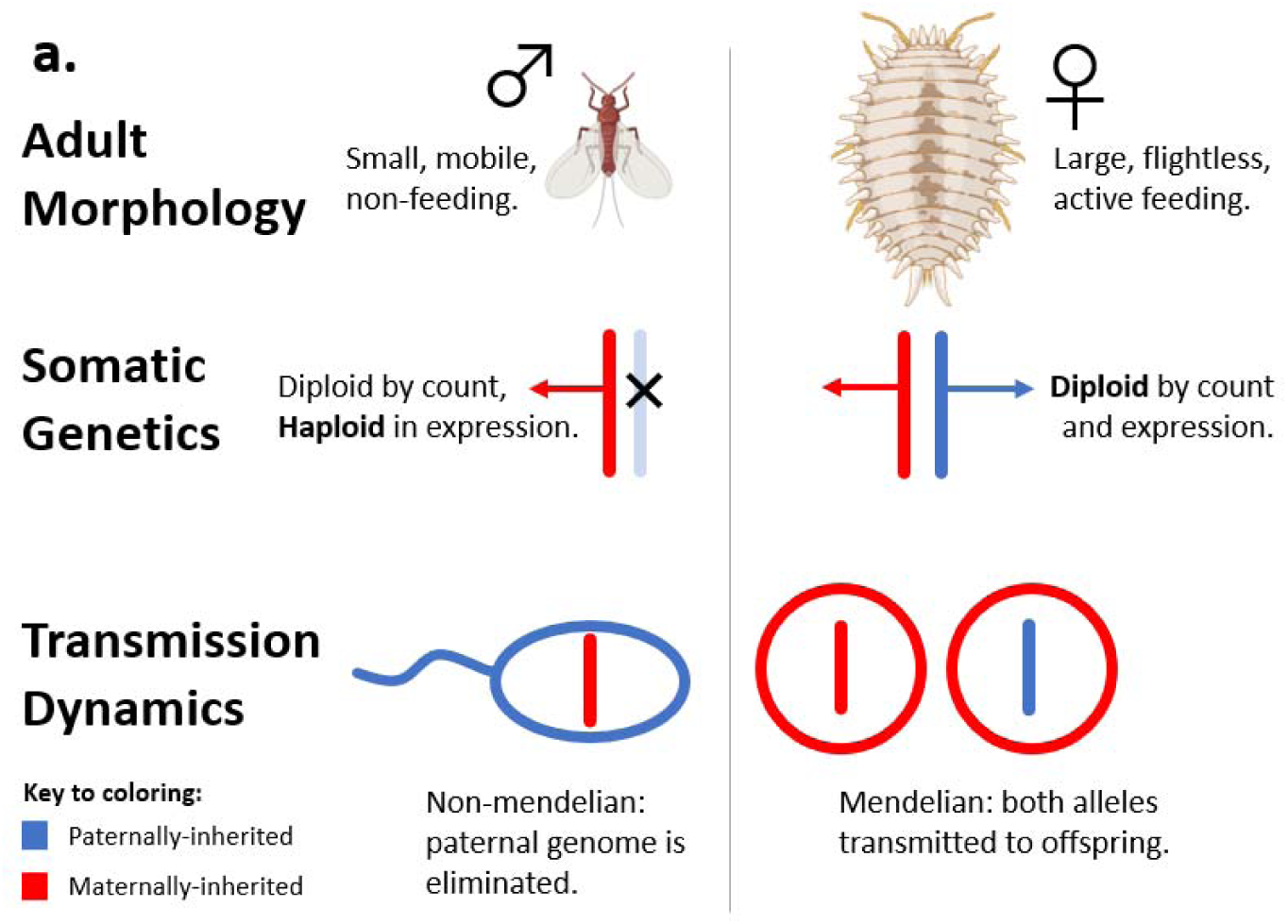
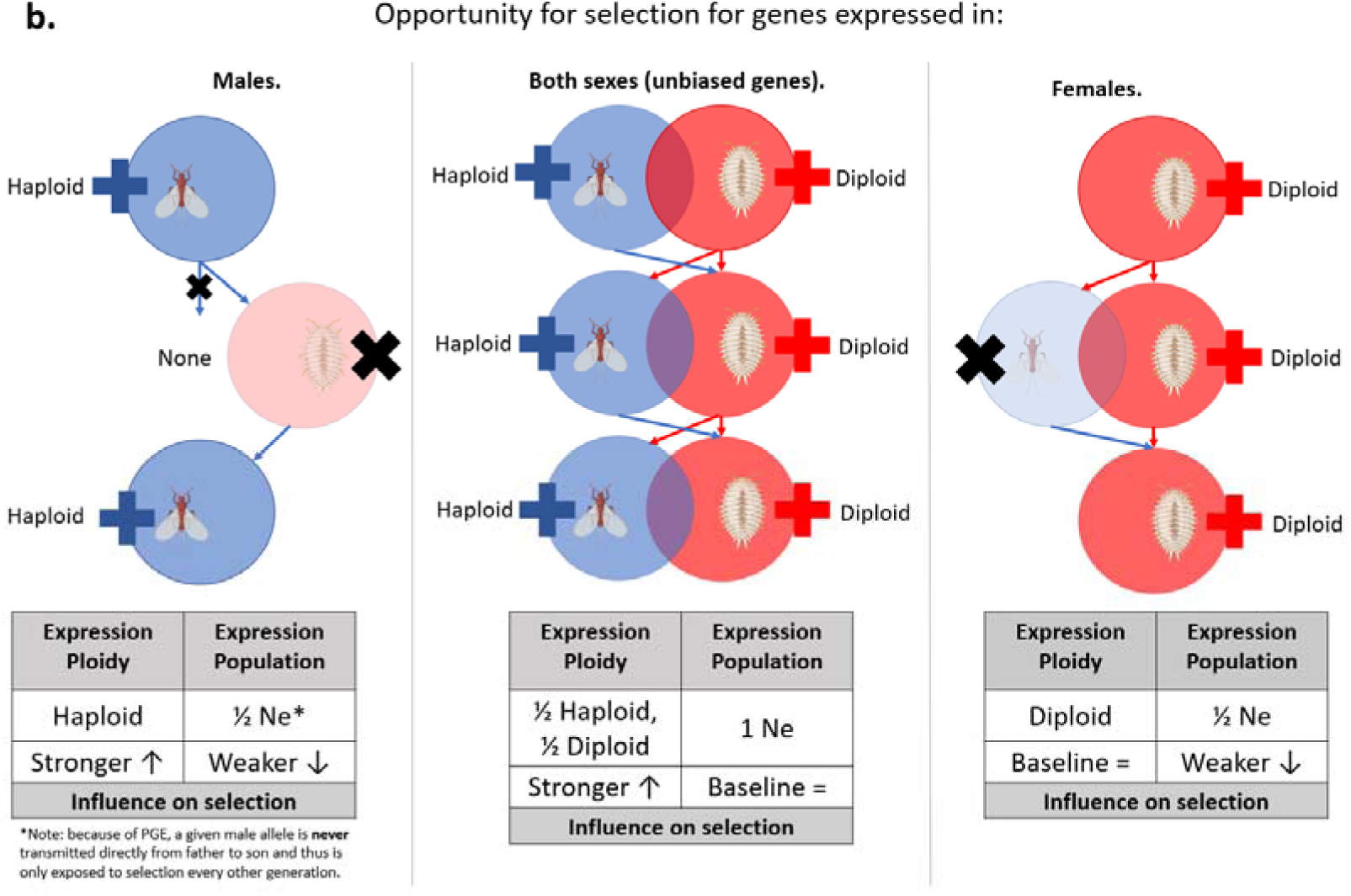
Introduction to the sex determination system of citrus mealybugs and the conditions it creates for sex-biased genes. **a. Overview of paternal genome elimination (PGE).** Males and females have extremely different morphologies despite a lack of sex chromosomes. Throughout their lives males only express one copy of their genome, their maternal haplotype. At time of reproduction, only maternal alleles are passed to the next generation by males. Females express and pass on both maternal and paternal alleles. **Considerationsb. Considerations for selection on sex-biased genes under PGE**. Alleles are exposed to selection (i.e., expressed) under very different conditions depending on the sex of expression. Male-biased genes are only expressed in half the population under equal sex ratio, are haploid in expression, and must pass from father to daughter to be expressed in a male again in the following generation. Unbiased genes are expressed in the full population, but in a haploid or diploid state depending on whether they are in males or females, respectively. Finally, female-biased genes are also only expressed in half the population, and only in a diploid state. Representations of mealybugs were created with biorender.com.

Although the precise molecular mechanisms controlling PGE and the initial genomic changes that enabled it are still unknown, the outcome is simple from a population genetic point of view. The mealybug PGE system is ultimately a form of haplodiploidy. However, unlike the most prominent examples of haplodiploidy found in Hymenoptera, especially among eusocial insects (Crozier et al. 1987; Gadau et al. 2000), both sexes of mealybug derive from sexual reproduction. Selective pressures are not complicated by complex social castes that create multiple distinct phenotypes for one sex, and reproduction is not restricted to a small subset of the population.

To a first approximation, PGE makes the entire mealybug genome similar to an X sex chromosome in some senses: it is expressed in the haploid state in males and spends more time over the generations in females than males (Hitchcock et al. 2022). But there are significant differences that make population genetic study of PGE an important contrast to the study of sex chromosomes. First, the sex chromosomes are predicted (Vicoso and Charlesworth 2009) and often observed (Mank et al. 2010; Mongue et al. 2022) to have a smaller effective population size than the autosomes. Second, owing to sex-limited recombination in some lineages, the sex chromosomes often have different effective recombination rates compared to the autosomes (Turner and Sheppard 1975; John et al. 2016). Third, sex chromosomes often evolve expression regulation systems to compensate for imbalances in copy number between the autosomes and sex chromosomes in the haploid sex (Disteche 2012; Gu et al. 2019). In each of the above sex chromosome scenarios, study of sex-biased genes is complicated by the fact that the sex chromosomes evolve under different dynamics and typically hold proportionally more sex-biased genes than the autosomes (Allen et al. 2013; Mongue and Walters 2017; Mongue and Baird 2024). The whole-genome nature of PGE ensures that sex-biased genes have the same background evolutionary dynamic regardless of linkage. For instance, recombination only occurs in female mealybugs (Bongiorni et al. 2004), but because PGE applies to the whole genome, recombination rates are not biased towards one part of the genome with more sex-biased genes.

To explore this unique form of sex determination, we study how sex-biased genes evolve in PGE, which creates differing and contradictory selective pressures on genes depending on their sex of expression (Figure 1b). In particular we ask (1) do genes with sex-biased expression generally evolve differently than those expressed in both sexes? (2) does haploid expression in males create more efficient selection? (3) how do these two factors interact to create the overall selective dynamic of paternal genome evolution at different evolutionary timescales?

## Methods

### Study system and sample collection

We studied the easily cultivated (Mahmoud et al. 2017) and widely invasive citrus mealybug, *Planococcus citri*. We reared colonies to generate RNA sequencing for nymphal and adult males and females well as proteomic data from the bacteriome and residual body tissue. For the RNA-Seq, mealybug colonies (strain CP1-2) were kept in a temperature and light controlled room at 25⁰C with a 16:8 light:dark photoperiod. They were fed *ad libitum* on sprouted Albert Bartlett Anya seed potatoes. Males and females were separated before sexual maturity to ensure virginity. Male and female nymphs were collected 18 days post-egg laying. Adult males were collected daily upon eclosion (aged 24-28 days) and adult females were collected between 32-35 days old. RNA was extracted from 30-50 3^rd^ instar males, 20 3^rd^ instar females, 30-50 adult males and 3 adult females per replicate, following a custom protocol (https://github.com/agdelafilia/wet_lab). We have previously shown that this does not affect downstream analysis (Bain et al. 2021). RNA quantity and quality were checked using Nanodrop and Qubit fluorometers as well as via gel electrophoresis. Samples were sent to BGI Tech Solution Co., Ltd. (Hong Kong) for library preparation and sequencing. Samples were sequenced to a depth of 50million reads per sample on a DNBSEQ platform using 150bp paired-end reads. We used these above data to provide evidence for gene annotation and, in the case of the RNAseq, establish the sex-bias in gene expression, as described below.

To understand the molecular evolution in nature, we sampled wild-caught individuals. We collected and sequenced one adult female per tree across a transect of citrus grove in Portugal. We extracted DNA from whole body tissues via a simple salt and alcohol precipitation and sequenced on an Illumina Novaseq to roughly 20x depth with Novogene (Cambridge, UK). See data availability for accession numbers.

We worked with the Darwin Tree of Life initiative to generate a high-quality genome assembly from a lab-reared colony of *P. citri* (Laura Ross et al. 2024). In brief, they used a mixture of PacBio HiFi long reads and Hi-C linked Illumina reads to generate an assembly that contains 5 chromosomal scaffolds. This external group did not do in-depth gene annotation, so we annotated the final genome with the BRAKER2 pipeline (Brůna et al. 2021), using as evidence protein sequences from the related mealybug *Phenacoccus solenopsis* (Li et al. 2020), proteins from a previous annotation of *P. citri* that were supported by mass spectrometry data, and newly generated RNAseq from nymphs and adults described directly below.

### Differential gene expression

Our expression dataset contained 4 biological replicates of third instar males (the earliest stage as which sex is morphologically distinguishable), 6 third instar females, 3 adult males, and 6 adult females; the smaller size of males results in higher failure rates of extraction and library prep, resulting in the uneven sampling between sexes. We aligned these expression datasets to the reference and annotation generated above and used RSEM v.1.3.3 (Li and Dewey 2011) implementing STAR v.2.7.10a (Dobin et al 2013) to quantify expression. We then used the R package DESeq2 v.1.40.1 (Love et al 2014) to identify differentially expressed genes between comparisons, using a Log_2_ fold-change > 1.5 and an adjusted p-value < 0.05. Specifically, we completed pairwise contrasts for male vs female expression in nymphs and in adults separately. We classified genes that passed our expression cutoffs but did not show significant sex-bias in expression as unbiased in subsequent analyses.

Not combining data for nymphs and adults gave us the opportunity to explore the consistency of sex-biased gene expression across life-stages. Because our expectations for molecular evolution are based on the ploidy of expression, such consistency is meaningful. For instance, a gene identified as significantly male-biased in adults, but expressed in both sexes in nymphs likely faces a different exposure to selection than one consistently male-biased in both juveniles and adults. We grouped genes into the categories shown in Table 1 based on significant results in nymphs and adults respectively. In the main analyses, we focus on the subset of genes with consistent bias in juveniles and adults (bolded diagonal), because our intention is to study the effects of PGE and ploidy-differences throughout life. For completeness, we also explore the small subset of genes that partially escape silencing in males (de la Filia et al. 2021), but finding no meaningful patterns, we report it in the supplement.

**Table 1.** Assignment of sex-biased genes based on expression profiles in both nymphal and adult mealybugs. We completed differential expression analyses between the sexes for nymphs and adults separately, then compared results. Categories along the diagonal showed consistent expression patterns, while those off-diagonal disagreed between nymphs and adults. We focus analyses on those with consistent expression (bold above) in the main text.

|  | Adult male-biased | Adult unbiased | Adult female-biased |
| --- | --- | --- | --- |
| Nymph male-biased | **Male-biased** | Partially male-biased | Sex reversal |
| Nymph unbiased | Partially male-biased | **Unbiased** | Partially female-biased |
| Nymph female-biased | Sex reversal | Partially female-biased | **Female-biased** |

### Population genomic analysis pipeline

We received adapter- and quality-trimmed reads from the sequencing company, making them alignment-ready. We took these data through a pipeline established by Mongue et al. (2019, 2022a; 2022b) to generate high quality SNV (single nucleotide variant) calls. We generated both within population (polymorphism) and between species (divergence) variant calls. For the latter, we used a previously-sequenced sample (de la Filia et al. 2021) of the related mealybug, *Planococcus ficus*. The process is described in detail below.

### Polymorphism

We aligned conspecific reads to the reference with Bowtie2 version 2.3.5.1’s *very-sensitive-local* alignment settings (Langmead and Salzberg 2012), sorted alignments and removed optical duplicate reads with Picard tools version 2.18.7 (Wysoker et al. 2013), then took these alignments through the Genome Analysis Toolkit version 4.1.9.0’s best practices pipeline for generating high-quality SNP calls (McKenna et al. 2010). Lacking a set of “known” SNPs for *P. citri*, we implemented a hard quality filter with the following parameters: Quality by Depth > 2.0, Fisher Strand-bias < 60, and Mapping Quality > 40. We continued analyses with SNPs that passed all of these filters.

Next, we used our newly generated gene annotations to create a custom database in the program SnpEff version 5.1d (Cingolani et al. 2012). This database parsed coding sequences into codons and classified all SNPs by their predicted amino acid effect (synonymous, missense, nonsense, etc.). To deal with cases of multiple transcripts belonging to a single gene, we used the -canon option to select the longest transcript as the canonical one. We took the subset of SNPs labeled “synonymous” to use as synonymous variants and considered only “missense” SNPs as non-synonymous under the logic that frameshifts and changes to start and stop codons are likely to cause large fitness effects across a gene and violate assumptions of SNP-based tests of selection.

Finally, some downstream tests for adaptation require information about the frequency of derived (new mutant) alleles compared to their ancestral state. Standard variant call format files (vcfs) contain estimates of *non-reference* allele frequency (AF), i.e., those that differ from the reference sequence. In the case that the reference allele is ancestral, then this frequency is the same as the derived allele frequency. If, however, the reference genome carries a derived allele, then AF represents the frequency of the ancestral allele instead. This is easily corrected by taking the complement (1 – AF) but requires knowledge of which alleles are ancestral and which are derived. We compared our polymorphism dataset with the divergence dataset below to infer ancestral allele state via parsimony as follows. If a position in the genome had a variant in the polymorphism dataset but not the divergence data, it meant that the outgroup carried the reference allele, and we concluded that the non-reference allele was derived. On the other hand, if polymorphism and divergence dataset shared the position and identity of an allele (ancestral polymorphisms in the section below), we inferred that the reference allele was ancestral and corrected the allele frequency to 1 – AF.

With these corrections completed, we tallied counts of non-synonymous polymorphisms (Pn) and synonymous polymorphisms (Ps) for each gene, both overall (for all variant frequencies) and using a sliding cutoff which excluded variants with a derived allele frequency < X where X ranged from 0.1 to 0.9 in increments of 0.1, using scripts previously developed and published by Mongue and Baird (2024). We also used an R script developed and published in Mongue et al., (2019) to label the degeneracy of each coding site position and sum for each gene to give us the number of non-synonymous and synonymous sites for each gene. With these values, we could normalize variant counts by the number of sites within a given gene. We combined these values to calculate pN/pS, or the scaled non-synonymous polymorphism rate. This value is the conceptual equivalent of dN/dS within species and measures the rate of non-synonymous polymorphism relative to synonymous polymorphism with each of the two categories normalized by the number of nonsynonymous or synonymous sites in a gene, respectively.

We tested between the different classes of genes (i.e., male-biased, unbiased, female-biased) using the non-parametric equivalent of an ANOVA, the Kruskal-Wallis test, first for pN (nonsynonymous polymorphisms per nonsynonymous site), then pS (synonymous polymorphisms per synonymous site), and finally for pN/pS together. For significant results, we performed a post-hoc Dunn’s test using the dunnTest() function in the R package “FSA” (Ogle and Ogle 2017) and used a threshold of p < 0.05 on Holm-Bonferroni adjusted p-values to determine significance of pairwise differences.

### Divergence

To obtain high-quality divergences (substitutions between species), we used a very similar pipeline to the one for polymorphism data. The starting point sequencing came from a single female *Planococcus ficus*, previously sequenced (de la Filia et al. 2021). Thanks to the relatively recent divergence time between *P. ficus* and *P. citri*, we were able to use the much more developed resources of the latter to simplify analyses. We aligned this outgroup to the *P. citri* genome again using bowtie2 and an identical set of downstream filtering parameters to generate and annotate a set of high-quality single nucleotide variants (in this case, divergences). We used the *P. citri* annotation to categorize these variants as synonymous or nonsynonymous, then sent them through an additional curation step to remove false divergences.

Aside from the methodological convenience, the similarity between the two species created a concern that ancestral polymorphism shared by both species could be included in naïve Dn and Ds counts. To remove these problem variants, we curated our dataset using the following logic. For the 330,360 homozygous coding sequence variants, we checked whether they shared both position (scaffold and base number) and identity (A,G,C,T) of a variant in the polymorphism dataset (e.g., REF = A, Focal.ALT = T, outgroup.ALT = T). If so, we marked these variants as ancestral to the split between species (ancestral polymorphism) and removed them from our count of divergences. If either the position or identity differed between outgroup and focal variants, we considered the divergence valid. Ultimately, 91.8% of homozygous variants passed this curation step.

For heterozygous variants, we again first parsed whether or not the outgroup variant intersected a focal variant. If not, then we considered the variant identity. Specifically, if a heterozygous outgroup variant was called as a single base (e.g. REF = A, ALT = T), this implied that the outgroup shared a reference allele (so the full genotype of the previous example would be A,T), and we removed these variants as false-divergences. In some cases however, the heterozygous outgroup site had two allele calls (e.g., REF = A, ALT = T,C). These were far rarer but could represent cases of a true divergence between species followed by a polymorphism at the same site. As a practical matter however, we had no way of assessing which variant was the divergence and which the polymorphism, so we kept only sites for which the two outgroup alleles had the same codon effect (synonymous or non-synonymous). In that way we could count the category of divergence without having to confidently determine to which of the two variants it belonged. In total there were only 411 of these tri-allelic true variants in our dataset.

Finally, we considered heterozygous outgroup variants that did coincide with focal polymorphisms. In these cases, the outgroup had to contain neither the reference allele nor the focal polymorphism(s) to be considered a true divergence. In other words, the only passing variants in this category were quadrallelic sites, which are vanishingly rare. In total, we found two valid 2 variants out of our 23,770 heterozygous coding sequence variants. Combined with preceding class of tri-allelic passing variants, only 1.7% of heterozygous outgroup variants represented true divergences. Because homozygous variants were far more common however, a total of 85.7% of our overall outgroup variants passed this curation step, leaving us with 303,981 coding sequence divergences for use in downstream analyses. As with the polymorphism data, we used the non-parametric Kruskal-Wallis test and post-hoc Dunn’s test to determine which groups (dN, dS, and dN/dS) were different from each other on a pairwise basis.

### Adaptation

Finally, we combined polymorphism and divergence data to estimate adaptive molecular evolution. We computed the proportion of amino acid substitutions driven by positive selection, α, in multiple ways. First, we used a simple-per gene calculation (Smith and Eyre-Walker 2002): α = 1 – (Pn/Ps)/(Dn/Ds). This statistic assumes that polymorphisms should represent mostly neutral variation, but in practice many populations show an excess of polymorphisms often attributed to a wealth of weakly deleterious polymorphisms that have not yet been removed from the population by selection (Charlesworth and Eyre-Walker 2008); this dynamic can result in negative α values which are not strictly defined and obscure the true proportion of adaptive substitution. These weakly deleterious variants should be almost exclusively nonsynonymous, as selection should only act directly on synonymous variants in rare cases of biased codon usage (Hershberg and Petrov 2008). To explore the extent of weakly deleterious variants in our dataset, we first plotted scaled nonsynonymous polymorphisms for each sex-bias class (female-biased, unbiased, and male-biased) of genes as a function of their derived allele frequency (Figure S1). Based on evidence of excess polymorphism at frequencies <0.2 in all gene classes, we chose this as cutoff and recalculated α = 1 – (Pn>0.2/Ps)/(Dn/Ds). This approach is similar to the heuristic used by Charlesworth and Eyre-Walker (2008), who employed a cutoff of Pn >0.15 to exclude weakly deleterious polymorphisms. We present the results of these two calculations (simple α and α removing low-frequency polymorphisms) in the main text. For added confidence in the robustness of the pattern, we explore a stricter cutoff, removing Pn < 0.4, in the supplement. These α calculations are compound statistics involving ratios and small count data, so, as with other population genetic statistics, used non-parametric statistics to assess significant differences using a Kruskal-Wallis test and post-hoc Dunn’s test if the former indicated significant differences.

### Long-term evolution: orthology

In the above analyses, we saw significant differences in molecular evolution of sex-biased gene classes in the short-term (polymorphisms within a *P. citri* population) and over the medium-term (divergence and adaptation between two *Planococcus* species). To explore these patterns across deeper evolutionary time, we compared evolutionary conservation of sex-biased genes across scale insect families. There are relatively few genomic resources for scale insects, but we found a high quality gene annotation for the Chinese wax scale, *Ericerus pela* (Yang et al. 2019), a member of the family Coccidae, which last shared a common ancestor with *P. citri* (Pseudococcidae) >150mya in the Jurassic (Vea and Grimaldi 2016). We called 1-to-1 orthologs between the two species’ gene annotations using the tool proteinortho (Lechner et al. 2011) v5.16 and then used a X^2^ test of independence to determine whether different sex-biased classes, as defined in *P. citri*, were conserved at similar or different rates.

## Results

### Gene annotation

We annotated 21,273 genes encoding 22,918 proteins and benchmarked this annotation against the BUSCO hemiptera_odb10 dataset of conserved orthologs. We recovered C:95.0 % [S:77.9 %, D:17.1 %], F:0.5 %, M:4.5 %. We note that BUSCO duplication scores are strongly impacted by the number of alternative transcripts in an annotation, so we parsed out the longest transcript per annotated gene, and then reran the BUSCO search, which lowered the apparent duplication rate to 8.8 %. We do not include this trimmed annotation as a resource because it discards potentially meaningful information about alternatively spliced genes; we only used this as a means to get a more accurate sense of ortholog duplication in the genome. More generally, at either duplication rate, this new annotation is undoubtedly a marked improvement over the previous annotation of 40,620 genes based on a much more fragmented assembly (https://ensembl.mealybug.org/Planococcus_citri_pcitriv1/Info/Index).

### Differential gene expression

We assessed differential gene expression between males and females in nymphs and adults separately. Of the 21,273 genes in the annotation, we excluded 6,676 genes mainly for low or no expression, but also for discrepancies in gene model lengths in a small number of cases (i.e. the exonic length was not divisible by three). This left us with 14,597 genes to analyze. Next, we carried out differential expression analyses separately for nymphal males vs females and for adult males vs females, then cross-referenced the results from the two life stages. There was strong agreement between life-stages over sex-bias of expression. Where nymphal and adult data disagreed, it typically manifested as more sex-biased genes in adults than nymphs. Cases of reversal of sex-bias (e.g. female-biased in nymphs but male-biased in adults) were a small fraction of expressed genes (Table 2). For the remainder of the analyses we focused on 7,322 genes with a consistent sex-bias in expression: 1003 male-biased (13.7%), 5155 unbiased (70.4%), and 1164 female-biased genes (15.9%). Although this doubtless excludes many genes in the genome, it represents the set of genes for which we have the highest confidence in expression bias and gives us large samples with which to make comparisons between bias classes.

**Table 2.**
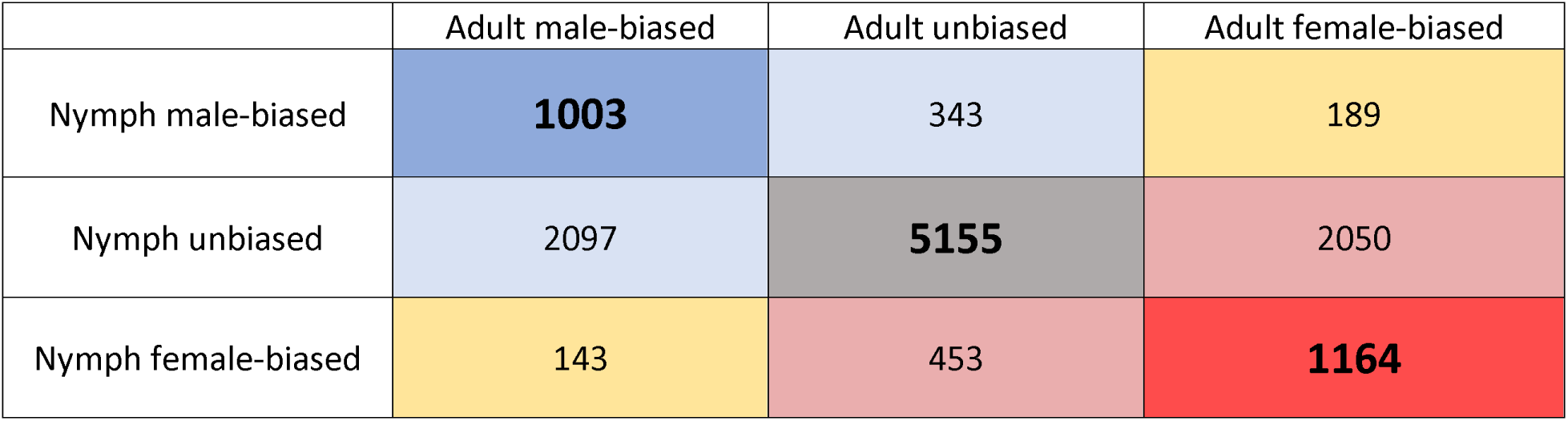
Categorization of sex-biased genes from differential expression analyses. We performed two separate but equivalent differential expression analyses to determine whether a given gene showed significant sex bias in expression: one using nymphal RNAseq and one using adult data. Values along the diagonal showed agreement in both tests. Unbiased genes are those expressed in both sexes equally. In the main text we focus on genes with consistent sex-bias in expression in both life stages (bold).

|  | Adult male-biased | Adult unbiased | Adult female-biased |
| --- | --- | --- | --- |
| Nymph male-biased | **1003** | 343 | 189 |
| Nymph unbiased | 2097 | **5155** | 2050 |
| Nymph female-biased | 143 | 453 | **1164** |

### Short-term evolution: sex differences in polymorphism

Using the above definitions of sex-biased genes, we compared polymorphism across the three consistently biased expression classes. We first examined nonsynonymous (i.e., putatively non-neutral) variation and synonymous (i.e., putatively neutral) variation separately, before combining the variation data to test the relative rate of nonsynonymous to synonymous variation within species (pN/pS).

For nonsynonymous variation, we found strong differences between the classes (X^2^ = 234.57, p < 0.0001). Post-hoc testing revealed the highest rates of scaled non-synonymous variation in female-biased genes, followed by male-biased genes, with unbiased genes holding the least (Table 3 Top, Figure 2 Top Left). Considering synonymous polymorphism alone, the sex-bias classes again differed from each other (X^2^ = 44.74, p < 0.0001) in the same pattern as nonsynonymous polymorphism (Table 3 middle and Figure 2 top right). Finally, combining pN and pS into the ratio pN/pS, we found a strong overall effect of sex-bias on the ratio of nonsynonymous to synonymous variation (X^2^ = 148.43, p < 0.0001).

**Figure 2.**
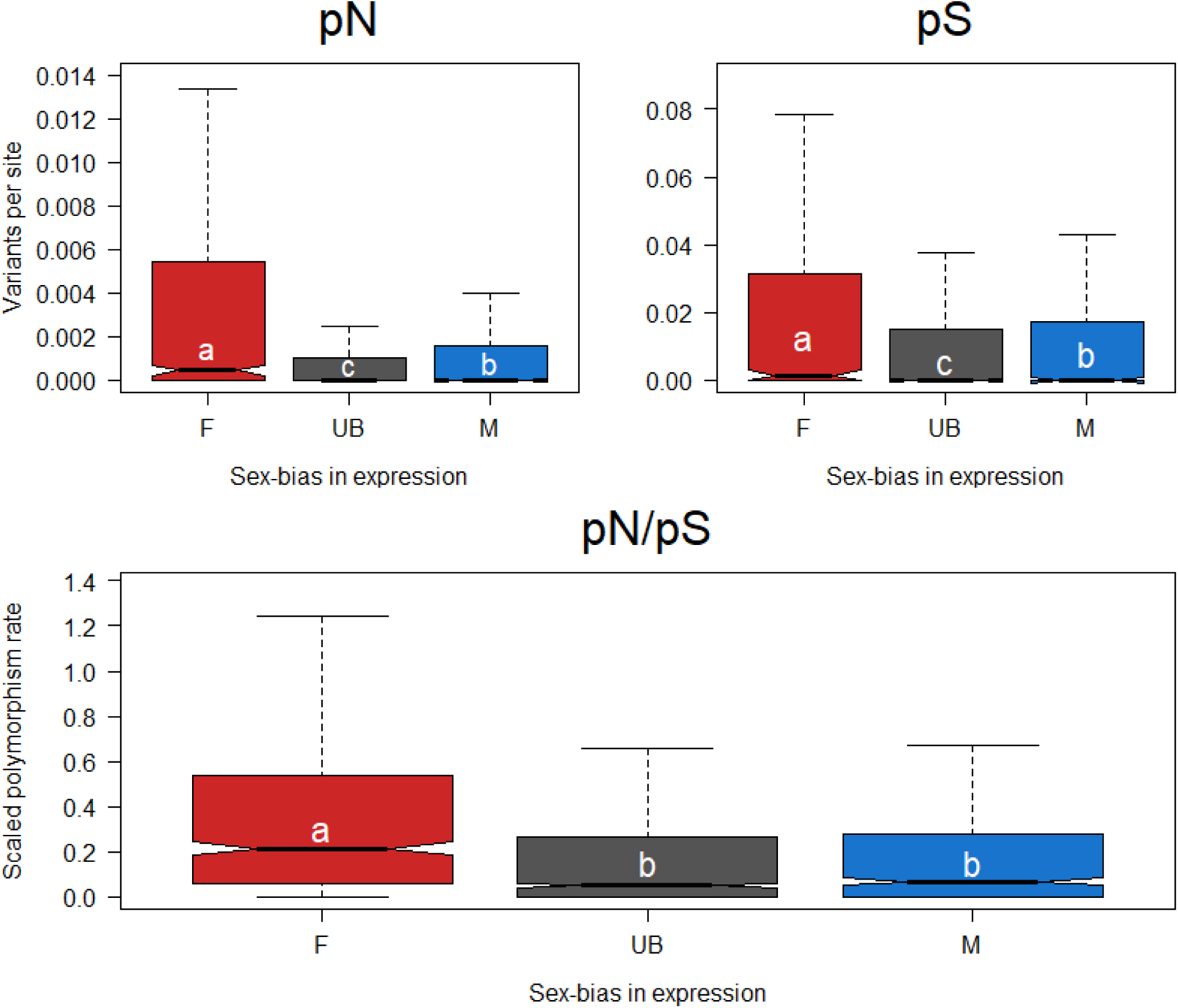
Polymorphism and sex-biased gene expression. Colors correspond to the bias classes defined in Table 1 and letters denote significant differences such that classes with different letters are significantly different and a > b > c. **Top left:** Nonsynonymous variants per nonsynonymous site (pN). Female-biased genes hold the most variation, followed by male-biased genes, then unbiased genes hold the least. **Top right:** Synonymous variants per synonymous site (pS). Synonymous variation follows the same pattern as nonsynonymous variation: female-biased genes hold the most, followed by male-biased and the unbiased genes. **Bottom:** Scaled polymorphism rate (pN/pS). Considering both classes together, female-biased genes hold more scaled variation than either unbiased ore male-biased genes, which do not differ from each other.

**Table 3.** Holm-Bonferroni adjusted p-values for pairwise differences between sex bias classes for within species variation (polymorphisms). Bolded values are significant at a p < 0.05 threshold. Top row: nonsynonymous variants per nonsynonymous site (pN). Middle: synonymous polymorphisms per synonymous site (pS). Bottom: scaled polymorphism (pN/pS).

| Statistic | Female-biased vs Unbiased | Unbiased vs Male-biased | Female-biased vs Male-biased |
| --- | --- | --- | --- |
| pN | **<0.0001** | **0.004** | **<0.0001** |
| pS | **<0.0001** | **0.021** | **0.004** |
| pN/pS | **<0.0001** | 0.113 | **<0.0001** |

These differences followed the differences observed in pN, with female-biased genes holding relatively more nonsynonymous variation than unbiased or male-biased genes (Figure 2 bottom, Table 3 Bottom).

### Long-term evolution: sex differences in divergence

Again, we first considered nonsynonymous substitutions per nonsynonymous site (dN). Again, sex-biased gene classes evolved differently from each other (Χ^2^_2_ = 59.75, p < 0.00001). Once again female-biased genes held the most variation, but for dN, unbiased genes held intermediate, followed by male-biased genes with the lowest divergence (Figure 3 top left, Table 4 Top). Then we considered synonymous substitutions per synonymous site (dS), which also varied significantly by sex-bias class (Χ^2^ = 153.85, p < 0.00001). The pattern here differs substantially from that in pS, with unbiased genes holding the most synonymous substitutions and male- and female-biased genes holding significantly less (Figure 3 top right, Table 4 Middle). To explain this unexpected pattern, we characterized the proportion of genes showing zero synonymous substitutions: 17.7% female-biased genes had no synonymous substitutions, 8.1% of male-biased genes and only 4.1% of unbiased genes. And finally, we found strong differences in rates of divergence across the sex-bias classes based on a Kruskal-Wallis test (Χ^2^ = 509.11, p < 0.00001). For overall scaled divergence (dN/dS) female-biased genes evolve the fastest, followed by male-biased, then unbiased genes (Figure 3 bottom, Table 4 Bottom).

**Figure 3.**
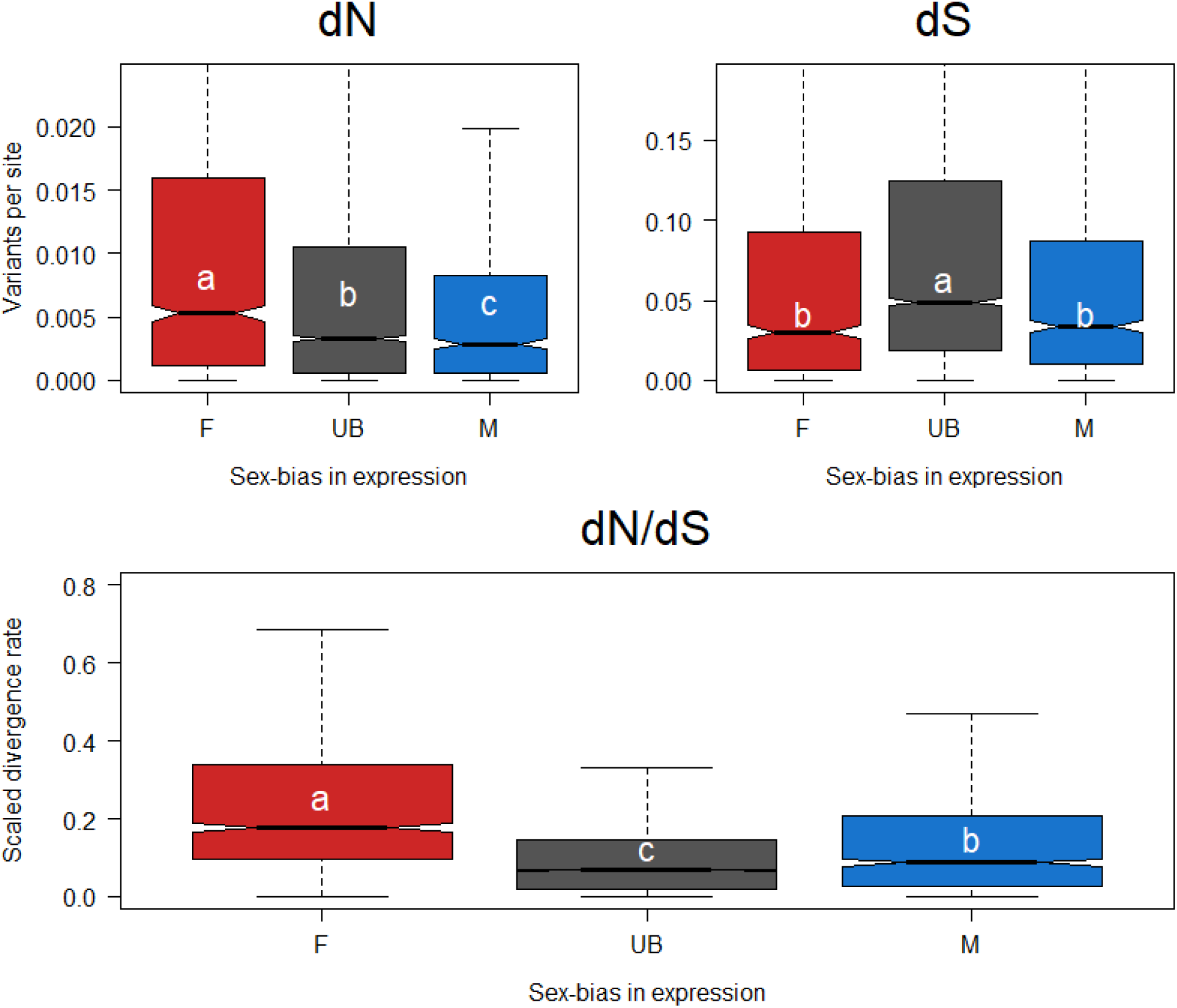
Divergence rates across sex-bias classes. **Top left:** nonsynonymous substitutions per non-synonymous site. Female-biased genes show the most nonsynonymous change, followed by unbiased genes, with male-biased genes show the least nonsynonymous change. **Top right:** Synonymous substitutions per synonymous site. Female-biased and male-biased genes show less scaled synonymous divergence than unbiased genes. **Bottom:** Scaled divergence (dN/dS). Overall female-biased genes evolve the fastest between species, followed by male-biased, and finally unbiased genes. Groups with different letters are significantly different from each other with values a > b > c, and colors follow the categorization from the methods.

**Table 4.** Holm-Bonferroni adjusted p-values for pairwise differences between sex bias classes for between species variation (divergences). Bolded values are significant at a p < 0.05 threshold. Top row: nonsynonymous substitutions per nonsynonymous site (dN). Middle: synonymous substitutions per synonymous site (dS). Bottom: scaled divergence (dN/dS).

| Statistic | Female-biased vs Unbiased | Unbiased vs Male-biased | Female-biased vs Male-biased |
| --- | --- | --- | --- |
| dN | **<0.0001** | **0.018** | **<0.0001** |
| dS | **<0.0001** | 0.085 | **0.004** |
| dN/dS | **<0.0001** | **<0.0001** | **<0.0001** |

### Adaptation across the mealybug genome

We calculated α first in a simple way, utilizing all polymorphism data from our focal population and found a significant difference between the sex-bias classes (Χ^2^_2_= 7.92, p = 0.019). Post-hoc testing revealed that this result is driven by lower adaptation in female-biased genes compared to unbiased genes (p = 0.016). Male-biased genes did not evolve strongly differently than female-biased genes (p = 0.093) or unbiased genes (p = 0.930, Figure 4 Left). We noted that α were very low, with many per-gene values being negative, a symptom of weakly deleterious polymorphisms sorting at low frequencies in the population; to account for this bias, we removed non-synonymous polymorphisms below 0.2 frequency in our dataset (Charlesworth and Eyre-Walker 2008) and recalculated α. With this approach, values more strongly differed (Χ^2^ = 21.54, p = 0.0002). Here, female-biased genes show less adaptation than both unbiased genes (p = 0.0.00001) or male-biased genes (p = 0.0036). Male-biased and unbiased genes do not evolve differently (p = 0.706, Figure 4 Right).

**Figure 4.**
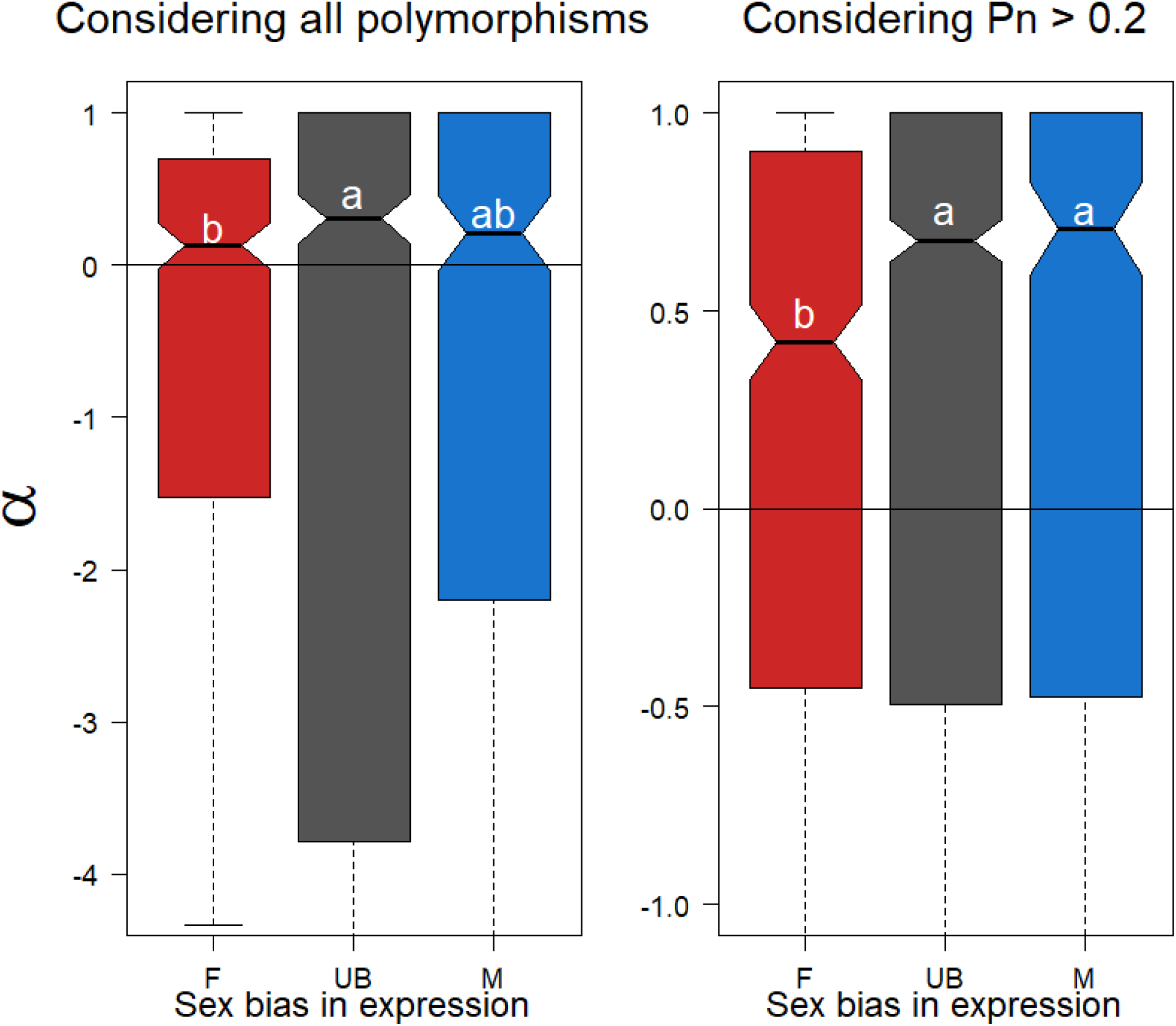
Adaptive evolution of sex-biased genes under PGE. The proportion of substitutions driven by positive selection (α) when considering all polymorphisms (left) or excluding non-synonymous polymorphisms with frequency < 0.2 (right). In both cases, female-biased genes show less adaptation than unbiased genes, although male-biased genes cannot be differentiated from either class when not filtering polymorphisms. After this filter however, both male-biased and unbiased genes evolve significantly more adaptively than female-biased genes. We checked the robustness of our findings by further increasing the stringency of to exclude non-synonymous polymorphisms < 0.4 frequency but found the same pattern of less adaptation in female-than unbiased or male-biased genes (Figure S2).

### Gene conservation across deeper evolutionary time

Of the 7,332 consistently sex-biased genes, we identified 2,878 with a 1-to-1 ortholog with the Chinese wax scale, *E. pela*. When parsing these orthologs by sex-bias in *P. citri*, we found significant difference between the classes (Χ^2^ = 842.94, p <0.0001). In particular, nearly half (2542, 49.3%) of unbiased genes showed strict conservation, compared to 259 (25.8%) of male-biased and only a mere 67 (5.7%) of female-biased genes.

## Discussion

### New genomic resources for the citrus mealybug in context with other mealybugs

We built upon the newly generated chromosomal assembly (Ross et al. 2024) to add value to the citrus mealybug as a model system. We generated 19 new RNAseq datasets, one proteomic dataset, and a set of gene annotations built on these new data. There are very few comparable sequencing efforts in mealybugs, but our annotations are consistent with recent resources generated for the obscure mealybug, *Pseudococcus viburni*, which has roughly 24,000 annotated protein coding genes (Vea et al. 2021) and stands in contrast to the annotation of the cotton mealybug, *Phenacoccus solenopsis*, with a mere 12,000 genes (Li et al. 2020). While the latter genome is roughly 30% smaller in total size than that of either *P. viburni* or *P. citri*, a near-two-fold difference in gene content is unlikely to be purely biological. And as both *P. citri* and *P. viburni* have BUSCO duplication rates <10 %, it is unlikely that the higher gene counts are substantially inflated by assembly artifacts. Instead, it is more likely that either the low number in *P. solenopsis* is an underestimate or the higher numbers reflect an increased rate of transposons annotated as genes in the species with larger genomes. Indeed, only around 15,000 genes were well-represented in our RNAseq, which may be closer to the true number of protein coding genes in *P. citri*, but with so few mealybug genomes available, it is currently difficult to generalize patterns of genomic natural history.

### Sexual dimorphism, paternal genome elimination, and mealybugs

With these resources established, we turn to evolutionary genetic questions, starting with the question of how much differential gene expression there is between the sexes. As expected based on gross morphology, citrus mealybugs are also highly sexually dimorphic at the gene expression level. Roughly one third of consistently expressed genes were significantly sex-biased and just over 60% were sex-biased in at least one life stage (adults or nymphs). Within this latter class, the vast majority of partially sex-biased genes show biased expression in adults but not juveniles. This observation fits well with both general predictions of sexual conflict theory and with the specific biology of mealybugs. In general, the fitness interests of both sexes are expected to be aligned in immature individuals as both sexes need to survive and grow; these interests diverge more sharply at sexual maturity when sex-specific reproductive strategies become relevant (Wedell et al. 2006). In mealybugs in particular, these differences extend far beyond the production of different gametes. Males undergo a process similar to metamorphosis which fundamentally changes their body plan with the addition of more developed sensory organs (e.g. eyes) and wings (Vea and Minakuchi 2021). Internally, adult males lose the endosymbiotic bacteria that provide juveniles of both sexes and adult females with essential nutrients (Kono et al. 2008). With so many internal and external differences, it is unsurprising that sexual dimorphism is strong at the gene expression level as well.

More specifically, recent theoretical work has predicted PGE should create a more favorable evolutionary dynamic for the invasion of female-biased alleles than male-biased, which over time should lead to feminization of gene expression (Hitchcock et al. 2022). And indeed, we see more female-biased genes than male-biased genes in *P. citri*. In terms of more familiar chromosomal sex determination, this places PGE’s expression profile closer to X chromosome systems (Prince et al. 2010; Perry et al. 2014) and sets it apart from Z chromosome species, which are often observed to have more male-biased than female-biased gene expression (Mongue et al. 2022; Mongue and Baird 2024). More to the point, the identification of sex-biased and unbiased genes allows us to answer the questions of molecular evolution that motivate this study.

### Short term variation: ploidy of expression creates sex differences

Both nonsynonymous and synonymous polymorphism individually followed the pattern of female-biased genes holding the most variation, followed by male-biased, then finally unbiased genes holding the least. Female-biased genes represent the more familiar diploid baseline from Mendelian genetic systems, so the overall results are best viewed as reductions in variation in unbiased and male-biased genes, both of which are exposed to haploid selection at least some of the time. This pattern matches observations that haploid expression in males strongly purges deleterious variants in other arthropods (Henter 2003; Tien et al. 2015) and confirms these expectation for PGE mealybugs (de la Filia et al. 2015). More surprisingly, this pattern extends to synonymous variants as well. As synonymous variants are rarely under direct selection (except, e.g., biased codon usage, Hershberg and Petrov 2008), this decrease in variation is likely the effect of background selection, which can remove neutral variants in linkage with sites under purifying selection (Charlesworth 2012). This possibility is supported by the observation that only one sex, females, have recombination in mealybugs, lowering the absolute rate of recombination compared to more familiar models (Bongiorni et al. 2004).

Parsing more finely, we saw even within haploid-expressed genes that male-biased genes hold more variation than unbiased genes. One traditional explanation for this pattern is that, being expressed in only half the population, male-biased genes should experience relaxed selection compared to unbiased genes (Gershoni and Pietrokovski 2014; Dapper and Wade 2016). While this general logic holds true, for male-biased genes under PGE, the reduced opportunity for selection is even more strict; any given male-biased gene currently expressed in a male will *not* be expressed in the next generation, needing to pass through a daughter in order to be expressed again in a grandson (de la Filia et al. 2015, 2021). As evidence for this reduced window of selection, we note that excess, likely deleterious polymorphisms reach higher frequencies (∼0.3) in male-biased genes than in either unbiased or female-biased genes (for which variation drops off above ∼0.1).

Finally, combining patterns of nonsynonymous and synonymous variants, we found that scaled polymorphism (pN/pS) was higher in female-biased genes, but unbiased and male-biased genes held similar ratios of variation. In summary, at the population level, PGE creates different patterns of variation based on the ploidy of expression. This result sets an expectation for how long-term variation should be distributed in the absence of positive selection (Kimura 1979; McDonald and Kreitman 1991; Ohta 1992).

### Long term evolution: sex-differences in speed and adaptation

We explored longer-term evolutionary differences first through patterns of divergence, fixed differences between *P. citri* and *P. ficus*. We found that female-biased genes had the highest scaled divergence (dN/dS), followed by male-biased genes, with unbiased genes evolving the slowest. This overall dynamic belies differences between how nonsynonymous and synonymous substitutions accrue, however. First, on the nonsynonymous side, the haploid-expressed genes evolve more slowly than female-biased genes, likely because of increased selective scrutiny. Furthermore, any particular male-biased allele can only be exposed to selection every other generation, which likely slows positive selection relative to unbiased genes, which can be expressed every generation. For synonymous divergence, the increase in substitutions in unbiased genes relative to female-biased genes may be the result of hitchhiking of neutral variation pulled to fixation when in linkage with a beneficial variant under positive selection (Maynard-Smith and Haigh 1974). This effect should be stronger in haploid-expressed genes where even recessive variation is under selection, but for male-biased alleles the requirement to pass through a daughter in between generations of selection likely gives more time for recombination to break up linkage between neutral and selected variants. Putting these patterns together, unbiased genes have the lowest dN/dS not because of a lack of nonsynonymous change, but rather because of a concomitant increase in synonymous divergence. More generally, the fact that relative rates of polymorphism and divergence differed between sex-bias classes suggests differing selective forces acting on these gene classes (McDonald and Kreitman 1991).

To directly examine adaptation, we estimated the proportion of adaptive substitutions, α, between *P. citri* and *P. ficus* for each class of genes. When considering all the polymorphisms in our dataset, we recovered fewer adaptive substitutions for female-biased genes than unbiased genes, but with male-biased genes indistinguishable from either group. When removing low-frequency, likely deleterious, polymorphisms that violate the assumptions of the α statistic (Charlesworth and Eyre-Walker 2008), we found that both unbiased and male-biased genes evolve more adaptively than female-biased genes. In other words, despite the predicted (Hitchcock et al. 2022) and observed feminization of the genome (this study), PGE ultimately creates conditions that favor adaptation for genes expressed in *males*.

Combined with the above results on dN/dS, this suggests that male-biased gene evolution is characterized by more adaptive, but slower change between species. To explore this dynamic over more distant evolutionary relationships, we compared conservation of 1-to-1 orthologs between *P. citri* and *Ericerus pela*, another PGE species from a separate hemipteran family, Coccidae (Yang et al. 2019). Using the sex-bias classifications from *P. citri*, we found the most conservation of unbiased genes, followed by male-biased genes, with female-biased genes in *P. citri* showing the fewest 1-to-1 orthologs between species, consistent with the pattern seen in *Planococcus* species.

### Comparison to chromosomal sex determination

In traditional chromosomal sex determination systems, sex-biased genes are often observed to evolve more adaptively on the sex chromosomes than autosomes (Meisel and Connallon 2013; Kousathanas et al. 2014; Sackton et al. 2014; Mongue et al. 2022), typically attributed to haploid selection on the X/Z in one sex. Conversely, when no increased adaptation is observed, the lower effective population size of the sex chromosome is often invoked to explain its absence (e.g., Mongue and Baird 2024). The PGE system is a useful contrast to the evolutionary dynamics of sex chromosomes for the simple reason that there is no tension between ploidy of expression and effective population size; these dynamics are consistent across the genome. Under these more straightforward conditions, we see that haploid expression of unbiased and male-biased genes increases efficacy of selection as predicted.

Less straightforward is the observation that some diploid-expressed genes do show increased adaptation on sex chromosomes compared to autosomes (e.g., female-biased X linked genes: Meisel and Connallon 2013; or male-biased Z-linked genes: Mongue et al. 2022). There is no ploidy benefit to being sex-linked in these cases. Instead, the simplest explanation is that, because of the biased transmission dynamics of sex chromosomes, sex-biased genes are more likely to find themselves in the correct sex for a favorable selective background if they become linked to a sex chromosome. This logic underpins the prediction (Klein et al. 2021) and observation that the X should accumulate female-biased genes and the Z should accumulate male-biased genes (e.g., Mongue and Walters 2017). Indeed, because of the transmission dynamics of PGE, the whole genome should favor invasion of female-biased alleles (Hitchcock et al. 2022). As a driver of adaptation, however, our data suggest it is far less powerful than haploid selection, because female-biased genes in *P. citri* show the lowest proportion of adaptive substitutions.

### Implications for pest management

Finally, mealybugs (Pseudococcidae) are the second largest family of scale insects (Hemiptera: Sternorrhyncha: Coccomorpha), with many species, including *P. citri*, being agricultural and ornamental pests of economic importance. A variety of control strategies have been developed, including chemical control (Franco et al. 2009) as well as more environmentally sustainable alternatives like mating disruption, mass-trapping, attract-and-kill, augmentative biological control, and classic biological control (Beltra et al. 2015; Sullivan et al. 2019; Franco et al. 2022; Gilliéron et al. 2024). Some of these tactics target one sex in particular, such as mating disruption targeting adult males or release of biological control parasitoids that seek female hosts. To be sure, more targeted studies of the evolutionary outcomes of mealybug pest management will be required, but our findings make testable predictions for the efficacy of different approaches. In particular, management that targets traits expressed in both sexes should be more likely to be effective across species, as the underlying genes appear more conserved over evolutionary time. Conversely, approaches that target only one sex, particularly females, may have more species-limited effects because of the faster divergence of female-biased genes.

However, given the evidence for less adaptation in these same genes, mealybug pests may be less likely to evolve resistance to female-targeted management practices. Integrating these molecular evolutionary considerations may help to develop robust control strategies with long term efficacy.

## Conclusions

We have generated resources and undertaken population genetic analyses of the citrus mealybug, *P. citri*, to better understand how sex-biased genes evolve under a non-chromosomal sex determination system. We found that (1) sex-biased genes, especially female-biased genes, evolve faster than genes expressed in both sexes; (2) haploid expression in unbiased and male-biased genes slows molecular change, likely from an increase in purifying selection; (3) in spite of the lower rate of change, the proportion of adaptive substitutions is higher for these unbiased and male-biased genes and (4) unbiased genes in particular are well-conserved across deep evolutionary time, with fewer male-biased and especially female-biased genes sharing a conserved ortholog with the soft scale, *E. pela*. Putting all of these observations together, we show that paternal genome elimination slows evolution but not adaptation in males, causing female-biased genes to vary more at multiple scales of evolutionary time. These results are consistent with theory developed in chromosomal sex determination systems, albeit without as many confounding factors that sometimes obscure them in those systems.

## Supporting information

Supplemental results

## Notes

### Competing Interest Statement

The authors have declared no competing interest.

## References

Albritton, S. E., A. L. Kranz, P. Rao, M. Kramer, C. Dieterich, and S. Ercan. 2014. Sex-biased gene expression and evolution of the X chromosome in nematodes. Genetics 197:865–883.

Allen, S. L., R. Bonduriansky, and S. F. Chenoweth. 2013. The genomic distribution of sex-biased genes in *Drosophila serrata*: X chromosome demasculinization, feminization, and hyperexpression in both sexes. Genome Biology and Evolution 5:1986–1994.

Arnqvist, G., and L. Rowe. 2005. Sexual Conflict. Princeton University Press.

Bain, S. A., H. Marshall, A. G. de la Filia, D. R. Laetsch, F. Husnik, and L. Ross. 2021. Sex-specific expression and DNA methylation in a species with extreme sexual dimorphism and paternal genome elimination. Molecular Ecology 30:5687–5703. Wiley Online Library.

Baines, J. F., S. A. Sawyer, D. L. Hartl, and J. Parsch. 2008. Effects of X-Linkage and Sex-Biased Gene Expression on the Rate of Adaptive Protein Evolution in Drosophila. Molecular Biology and Evolution 25:1639–1650.

Beltra, A., P. Addison, J. A. Avalos, D. Crochard, F. Garcia-Mari, E. Guerrieri, J. H. Giliomee, T. Malausa, C. Navarro-Campos, and F. Palero. 2015. Guiding classical biological control of an invasive mealybug using integrative taxonomy. PloS one 10:e0128685. Public Library of Science San Francisco, CA USA.

Blackmon, H., L. Ross, and D. Bachtrog. 2017. Sex determination, sex chromosomes, and karyotype evolution in insects. Journal of Heredity 108:78–93. Oxford University Press US.

Bongiorni, S., P. Fiorenzo, D. Pippoletti, and G. Prantera. 2004. Inverted meiosis and meiotic drive in mealybugs. Chromosoma 112:331–341. Springer.

Brůna, T., K. J. Hoff, A. Lomsadze, M. Stanke, and M. Borodovsky. 2021. BRAKER2: automatic eukaryotic genome annotation with GeneMark-EP+ and AUGUSTUS supported by a protein database. NAR genomics and bioinformatics 3:lqaa108. Oxford University Press.

Charlesworth, B. 2012. The role of background selection in shaping patterns of molecular evolution and variation: evidence from variability on the Drosophila X chromosome. Genetics 191:233–246.

Charlesworth, J., and A. Eyre-Walker. 2008. The McDonald–Kreitman test and slightly deleterious mutations. Molecular biology and evolution 25:1007–1015. Oxford University Press.

Cingolani, P., A. Platts, L. L. Wang, M. Coon, T. Nguyen, L. Wang, S. J. Land, X. Lu, and D. M. Ruden. 2012. A program for annotating and predicting the effects of single nucleotide polymorphisms, SnpEff: SNPs in the genome of *Drosophila melanogaster* strain w 1118; iso-2; iso-3. Fly 6:80–92.

Civetta, A., and R. S. Singh. 1995. High divergence of reproductive tract proteins and their association with postzygotic reproductive isolation in *Drosophila melanogaster* and *Drosophila virilis* group species. Journal of Molecular Evolution 41:1085–1095.

Crozier, R. H., B. H. Smith, and Y. C. Crozier. 1987. Relatedness and Population Structure of the Primitively Eusocial Bee Lasioglossum zephyrum (Hymenoptera: Halictidae) in Kansas. Evolution 41:902–910.

Danzmann, R. G., J. D. Norman, E. B. Rondeau, A. M. Messmer, M. P. Kent, S. Lien, O. Igboeli, M. D. Fast, and B. F. Koop. 2019. A genetic linkage map for the salmon louse (Lepeophtheirus salmonis): evidence for high male: female and inter-familial recombination rate differences. Molecular Genetics and Genomics 294:343–363. Springer.

Dapper, A. L., and M. J. Wade. 2016. The evolution of sperm competition genes: The effect of mating system on levels of genetic variation within and between species. Evolution 70:502–511.

de la Filia, A. G., S. A. Bain, and L. Ross. 2015. Haplodiploidy and the reproductive ecology of Arthropods. Current Opinion in Insect Science 9:36–43. Elsevier.

de la Filia, A. G., A. J. Mongue, J. Dorrens, H. Lemon, D. R. Laetsch, and L. Ross. 2021. Males that silence their father’s genes: genomic imprinting of a complete haploid genome. Molecular Biology and Evolution 38:2566–2581.

Dean, R., P. W. Harrison, A. E. Wright, F. Zimmer, and J. E. Mank. 2015. Positive selection underlies faster-Z evolution of gene expression in birds. Molecular biology and evolution 32:2646–2656.

Disteche, C. M. 2012. Dosage Compensation of the Sex Chromosomes. Annual Review of Genetics 46:537–560.

Ellegren, H. 2011. Sex-chromosome evolution: recent progress and the influence of male and female heterogamety. Genetics 12:157–66.

Franco, J.C., A. Cocco, A. Lucchi, Z. Mendel, P. Suma, S. Vacas, R. Mansour, and V. Navarro-Llopis. 2022. Scientific and technological developments in mating disruption of scale insects. Entomologia Generalis 42:251–273. Schweizerbart

Franco, J. C., A. Zada, and Z. Mendel. 2009. Novel approaches for the management of mealybug pests. Biorational control of arthropod pests: application and resistance management 233–278. Springer.

Gadau, J., R. E. Page Jr, J. H. Werren, and P. Schmid-Hempel. 2000. Genome Organization and Social Evolution in Hymenoptera. Naturwissenschaften 87:87–89.

Gavrilov, I. A. 2007. A catalog of chromosome numbers and genetic systems of scale insects (Homoptera: Coccinea) of the world. Israel Journal of Entomology 37:1–45.

Gershoni, M., and S. Pietrokovski. 2014. Reduced selection and accumulation of deleterious mutations in genes exclusively expressed in men. Nature Communications 5.

Gilliéron, F., J. Romeis, and J. Collatz. 2024. Cold tolerance of the mealybug parasitoid Anagyrus vladimiri. BioControl 69:129–143. Springer.

Gu, L., P. F. Reilly, J. J. Lewis, R. D. Reed, P. Andolfatto, and J. R. Walters. 2019. Dichotomy of Dosage Compensation along the Neo Z Chromosome of the Monarch Butterfly. Current Biology 29:4071–4077.e3.

Gu, L., and J. R. Walters. 2017. Evolution of Sex Chromosome Dosage Compensation in Animals: A Beautiful Theory, Undermined by Facts and Bedeviled by Details. Genome Biology and Evolution 9:2461–2476.

Henter, H. J. 2003. Inbreeding depression and haplodiploidy: experimental measures in a parasitoid and comparisons across diploid and haplodiploid insect taxa. Evolution 57:1793–1803. Blackwell Publishing Ltd Oxford, UK.

Herrick, G., and J. Seger. 1999. Imprinting and paternal genome elimination in insects. Pp. 41–71 *in* R. Ohlsson, ed. Springer-Verlag, New York:

Hershberg, R., and D. A. Petrov. 2008. Selection on codon bias. Annual review of genetics 42:287–299. Annual Reviews.

Hitchcock, T. J., A. Gardner, and L. Ross. 2022. Sexual antagonism in haplodiploids. Evolution 76:292–309. John Wiley & Sons, Ltd.

Hughes-Schrader, S. 1948. Cytology of coccids (Coccoidea-Homoptera). Adv. Genet. 35:127–203.

Jaquiéry, J., C. Rispe, D. Roze, F. Legeai, G. Le Trionnaire, S. Stoeckel, L. Mieuzet, C. Da Silva, J. Poulain, N. Prunier-Leterme, B. Ségurens, D. Tagu, and J.-C. Simon. 2013. Masculinization of the X Chromosome in the Pea Aphid. PLOS Genetics 9:e1003690.

John, A., K. Vinayan, and J. Varghese. 2016. Achiasmy: male fruit flies are not ready to mix. Frontiers in cell and developmental biology 4:75. Frontiers Media SA.

Kimura, M. 1979. The neutral theory of molecular evolution. Scientific American 241:98–129. JSTOR.

Klein, K., H. Kokko, and H. Ten Brink. 2021. Disentangling verbal arguments: intralocus sexual conflict in haplodiploids. The American Naturalist 198:678–693. The University of Chicago Press Chicago, IL.

Kono, M., R. Koga, M. Shimada, and T. Fukatsu. 2008. Infection Dynamics of Coexisting Beta- and Gammaproteobacteria in the Nested Endosymbiotic System of Mealybugs. Applied and Environmental Microbiology 74:4175–4184.

Kousathanas, A., D. L. Halligan, and P. D. Keightley. 2014. Faster-X adaptive protein evolution in house mice. Genetics 196:1131–1143.

Langmead, B., and S. L. Salzberg. 2012. Fast gapped-read alignment with Bowtie 2. Nat Methods 9:357– 359.

Laura Ross, Andrew J. Mongue, Andres de la Filia, Darwin Tree of Life Barcoding collective, Wellcome Sanger Institute Tree of Life Management, Samples and Laboratory Team, Wellcome Sanger Institute Scientific Operations: Sequencing Operations, Wellcome Sanger Institute Tree of Life Core Informatics team, Tree of Life Core Informatics collective, and Darwin Tree of Life Consortium. 2024. The genome sequence of the citrus mealybug, Planococcus citri (Risso, 1913). Wellcome Open Research 9.

Lechner, M., S. Findeiss, L. Steiner, M. Marz, P. F. Stadler, S. J. Prohaska, S. Findeiß, L. Steiner, M. Marz, P. F. Stadler, and S. J. Prohaska. 2011. Proteinortho: Detection of (Co-)orthologs in large-scale analysis. BMC Bioinformatics 12:124.

Li, M., H. Tong, S. Wang, W. Ye, Z. Li, M. A. A. Omar, Y. Ao, S. Ding, Z. Li, Y. Wang, C. Yin, X. Zhao, K. He, F. Liu, X. Chen, Y. Mei, J. R. Walters, M. Jiang, and F. Li. 2020. A chromosome-level genome assembly provides new insights into paternal genome elimination in the cotton mealybug Phenacoccus solenopsis. Molecular Ecology Resources n/a. John Wiley & Sons, Ltd.

Mahmoud, H. T., H. Nabil, A. Shahein, and Z. Mohamed. 2017. Biological studies on the citrus mealybug, planococcus citri (RISSO)(Hemiptera: Pseudococcidae) under laboratory conditions. Zagazig Journal of Agricultural Research 44:1097–1106. Zagazig University, Faculty of Agriculture.

Mank, J. E., K. Nam, and H. Ellegren. 2009. Faster-Z evolution is predominantly due to genetic drift. Molecular biology and evolution 27:661–670.

Mank, J. E., B. Vicoso, S. Berlin, and B. Charlesworth. 2010. Effective Population Size and the Faster-X Effect: Empirical Results and Their Interpretation. Evolution 64:663–674.

Maynard-Smith, J., and J. Haigh. 1974. The hitch-hiking effect of a favourable gene. Genet. Res. 23:23– 35.

McDonald, J. H., and M. Kreitman. 1991. Adaptive protein evolution at the Adh locus in Drosophila. Nature 351:652–4.

McKenna, A. H., M. Hanna, E. Banks, A. Sivachenko, K. Cibulskis, A. Kernytsky, K. Garimella, D. Altshuler, S. Gabriel, M. Daly, and M. Depristo. 2010. The Genome Analysis Toolkit: A MapReduce framework for analyzing next-generation DNA sequencing data. Genome Research 20:1297– 1303.

Meisel, R. P. 2011. Towards a more nuanced understanding of the relationship between sex-biased gene expression and rates of protein-coding sequence evolution. Molecular Biology and Evolution 28:1893–1900. Oxford University Press.

Meisel, R. P., and T. Connallon. 2013. The faster-X effect: integrating theory and data. Trends in Genetics 29:537–544.

Mongue, A. J., and R. B. Baird. 2024. Genetic drift drives faster-Z evolution in the salmon louse Lepeophtheirus salmonis. Evolution qpae090.

Mongue, A. J., M. E. Hansen, L. Gu, C. E. Sorenson, and J. R. Walters. 2019. Nonfertilizing sperm in Lepidoptera show little evidence for recurrent positive selection. Molecular Ecology 28:2517– 2530.

Mongue, A. J., M. E. Hansen, and J. R. Walters. 2022. Support for faster and more adaptive Z chromosome evolution in two divergent lepidopteran lineages *. Evolution 76:332–345.

Mongue, A. J., and A. Y. Kawahara. 2022. Population differentiation and structural variation in the *Manduca sexta* genome across the United States. G3 Genes|Genomes|Genetics 12:jkac047.

Mongue, A. J., and J. Walters. 2017. The Z chromosome is enriched for sperm proteins in two divergent species of Lepidoptera. Genome 61:248–253. NRC Research Press.

Nelson Rees, W. A. 1962. Effects of radiation damaged heterochromatic chromosomes on male fertility in mealy bug, Planococcus citri (risso). Genetics 47:661-amp;

Nur, U. 1980. Evolution of unusual chromosome systems in scale insects (Coccoidea: Homoptera). Insect cytogenetics 97–117. Blackwell Scientific Publications.

Nur, U. 1966. Nonreplication of heterochromatic chromosomes in a mealy bug Planococcus citri (Coccoidea - Homoptera). Chromosoma 19:439–448.

Ogle, D., and M. D. Ogle. 2017. Package ‘FSA.’ Cran Repos 1–206.

Ohta, T. 1992. The Nearly Neutral Theory of Molecular Evolution. Annual Review of Ecology and Systematics 23:263–286.

Perry, J. C., P. W. Harrison, and J. E. Mank. 2014. The ontogeny and evolution of sex-biased gene expression in Drosophila melanogaster. Molecular biology and evolution 31:1206–1219. Oxford University Press.

Prince, E. G., D. Kirkland, and J. P. Demuth. 2010. Hyperexpression of the X chromosome in both sexes results in extensive female bias of X-linked genes in the flour beetle. Genome biology and evolution 2:336–346. Oxford University Press.

Ross, L., A. J. Mongue, A. De La Filia, Darwin Tree of Life Barcoding collective, S. and L. team Wellcome Sanger Institute Tree of Life Management, Wellcome Sanger Institute Scientific Operations: Sequencing Operations, Wellcome Sanger Institute Tree of Life Core Informatics team, Tree of Life Core Informatics collective, and Darwin Tree of Life Consortium. 2024. The genome sequence of the citrus mealybug, Planococcus citri (Risso, 1913). Wellcome Open Research 9:22. F1000 Research Limited London, UK.

Ross, L., A. J. Mongue, C. N. Hodson, and T. Schwander. 2022. Asymmetric Inheritance: The Diversity and Evolution of Non-Mendelian Reproductive Strategies. Annu. Rev. Ecol. Evol. Syst. 53:1–23. Annual Reviews.

Rousselle, M., N. Faivre, M. Ballenghien, N. Galtier, and B. Nabholz. 2016. Hemizygosity enhances purifying selection: lack of fast-Z evolution in two satyrine butterflies. Genome biology and evolution 8:3108–3119.

Sackton, T. B., R. B. Corbett-Detig, J. Nagaraju, L. Vaishna, K. P. Arunkumar, and D. L. Hartl. 2014. Positive selection drives faster-Z evolution in silkmoths. Evolution 68:2331–2342.

Schrader, F. 1921. The chromosomes of Pseudococcus nipae. The Biological Bulletin 40:259–269. Marine Biological Laboratory.

Smith, N. G. C., and A. Eyre-Walker. 2002. Adaptive protein evolution in *Drosophila*. Nature 415:1022– 1024.

Sullivan, N., R. Butler, L. Salehi, A. Twidle, G. Baker, and D. Suckling. 2019. Deployment of the sex pheromone of Pseudococcus calceolariae (Hemiptera: Pseudococcidae) as a potential new tool for mass trapping in citrus in South Australia. New Zealand Entomologist 42:1–12. Taylor & Francis.

Swanson, W. J., and V. D. Vacquier. 2002. The rapid evolution of reproductive proteins. Genetics 3:137– 144.

Tien, N. S. H., M. W. Sabelis, and M. Egas. 2015. Inbreeding depression and purging in a haplodiploid: gender-related effects. Heredity 114:327–332.

Tree of Sex Consortium. 2014. Tree of Sex: A database of sexual systems. Scientific Data 1. Nature Publishing Group.

Turner, J. R. G., and P. M. Sheppard. 1975. Absence of Crossover in Female Butterflies (Heliconius). Heredity 34:265–269.

Vea, I. M., A. G. de la Filia, K. S. Jaron, A. J. Mongue, F. J. Ruiz-Ruano, S. E. Barlow, R. Nelson, and L. Ross. 2021. The B chromosome of Pseudococcus viburni: a selfish chromosome that exploits whole-genome meiotic drive. bioRxiv 2021–08. Cold Spring Harbor Laboratory.

Vea, I. M., and D. A. Grimaldi. 2016. Putting scales into evolutionary time: the divergence of major scale insect lineages (Hemiptera) predates the radiation of modern angiosperm hosts. Scientific Reports 6:23487. Nature Publishing Group UK London.

Vea, I. M., and C. Minakuchi. 2021. Atypical insects: molecular mechanisms of unusual life history strategies. Current Opinion in Insect Science 43:46–53.

Vicoso, B., and B. Charlesworth. 2009. Effective population size and the faster-X effect: an extended model. Evolution: International Journal of Organic Evolution 63:2413–2426.

Wedell, N., C. Kvarnemo, C. M. Lessells, and T. Tregenza. 2006. Sexual conflict and life histories. Animal Behaviour 71:999–1011.

Whittle, C. A., A. Kulkarni, and C. G. Extavour. 2020. Absence of a faster-X effect in beetles (Tribolium, Coleoptera). G3: Genes, Genomes, Genetics 10:1125–1136. G3: Genes, Genomes, Genetics.

Wright, A. E., P. W. Harrison, F. Zimmer, S. H. Montgomery, M. A. Pointer, and J. E. Mank. 2015. Variation in promiscuity and sexual selection drives avian rate of Faster-Z evolution. Molecular ecology 24:1218–1235.

Wright, A. E., T. F. Rogers, M. Fumagalli, C. R. Cooney, and J. E. Mank. 2019. Phenotypic sexual dimorphism is associated with genomic signatures of resolved sexual conflict. Molecular ecology 28:2860–2871. Wiley Online Library.

Wysoker, A., K. Tibbetts, and T. Fennell. 2013. Picard tools version 1.90. http://picard.sourceforge.net (Accessed 14 December 2016) 107:308.

Yang, P., S. Yu, J. Hao, W. Liu, Z. Zhao, Z. Zhu, T. Sun, X. Wang, and Q. Song. 2019. Genome sequence of the Chinese white wax scale insect Ericerus pela: the first draft genome for the Coccidae family of scale insects. GigaScience. 8:giz113. Oxford University Press.

