## Supplemental results for "Paternal genome elimination creates contrasting evolutionary trajectories in male and female citrus mealybugs"

**Supplementary analyses and results**

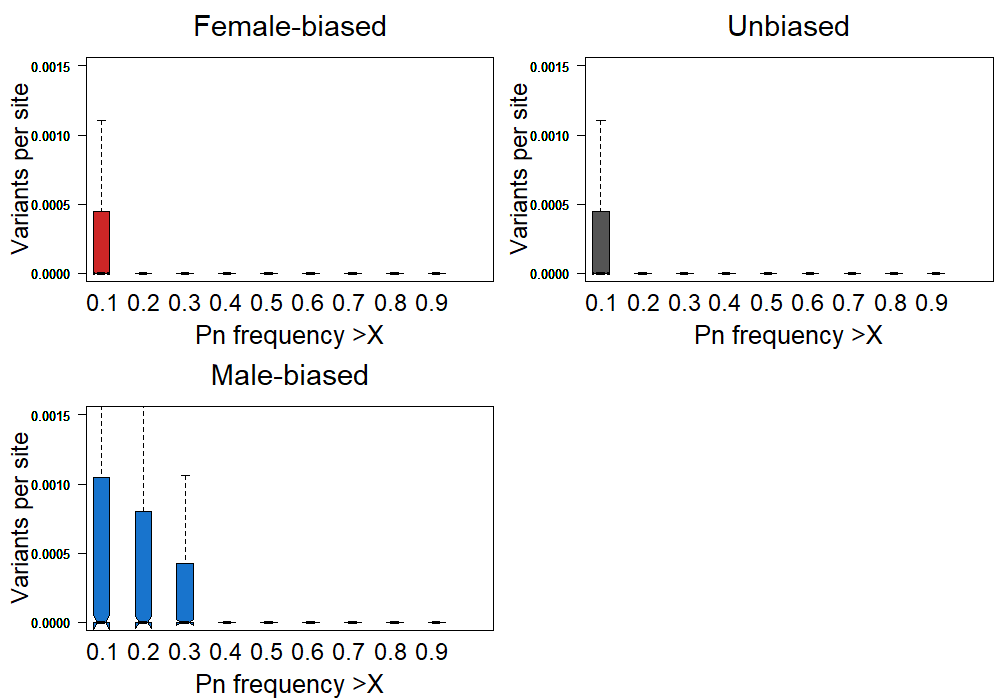

**Figure S1. Derived allele frequency of non-synonymous polymorphisms by sex-bias class.** All three gene classes show a massive excess of variation at low frequency (<0.2), suggesting a genome-wide phenomenon of weakly deleterious variation maintained in the population at low frequencies. For male-biased genes (bottom left) however, this excess of variation is maintained to much higher frequencies (<0.4), potentially due to the lack of direct transmission between fathers and sons, allowing these alleles to drift every other generation.

Based on this difference, we explored several cutoffs. First considering all polymorphisms or the more permissive removal of Pn < 0.2 (both reported in the main text) and finally the stricter removal of Pn < 0.4 (reported here). Again, we found a significant difference in α (Χ^2^_2_= 26.81, p < 0.0001), with female-biased genes evolving less adaptively than male-biased (p = 0.0013) or unbiased genes (p < 0.0001). Male-biased and unbiased genes do not evolve differently under these analyses ( p= 0.604; Figure S2).

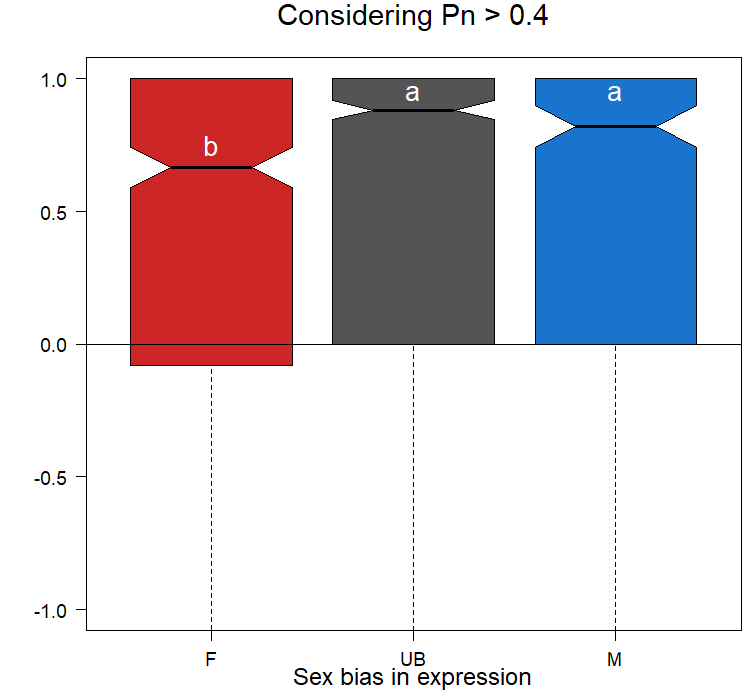

**Figure S2. Stricter filtering of putatively deleterious alleles.** Based on above results of excess polymorphism at higher frequencies in male-biased genes, we recalculated after excluding non-synonymous polymorphisms < 0.4 derived allele frequency. As with less strict filtering, male-biased and unbiased genes show similar α values, both of which are significantly higher than for female-biased genes.

*Investigation of escape of PGE*

Previous results from de la Filia et al. (2021) demonstrated that although PGE is a genome-wide phenomenon, it does not completely silence paternal alleles in males; there are a handful of genes that show some level of paternal expression. We reanalyzed this old dataset using our updated annotations to create a continuous variable, maternal expression, that ranged from 0 (completely paternal expression) to 1 (completely maternal expression, expected under PGE). By this metric, we recovered 20 male-biased genes and 188 unbiased genes with some level of paternal allelic expression. We tested whether these genes evolved differently via simple linear models relating the degree of maternal expression to dN/dS and α in unbiased and male-biased genes separately.

For unbiased genes, there was no significant relationship between maternal expression and dN/dS (F_1,52_ = 0.0005, p = 0.9823) or α, using the Pn >0.2 cutoff from the main text (F_1,53_ = 2.463, p = 0.1225). Likewise for male-biased genes, there was no significant relationship between maternal expression and dN/dS (F_1,9_ = 0.179, p = 0.682) or α (F_1,9_ = 3.918, p = 0.0792). These results are perhaps not surprising, as even for genes that escape strict paternal silencing, the vast majority still show biased expression, with the maternal allele making up > 50% of transcripts (Figure S3).

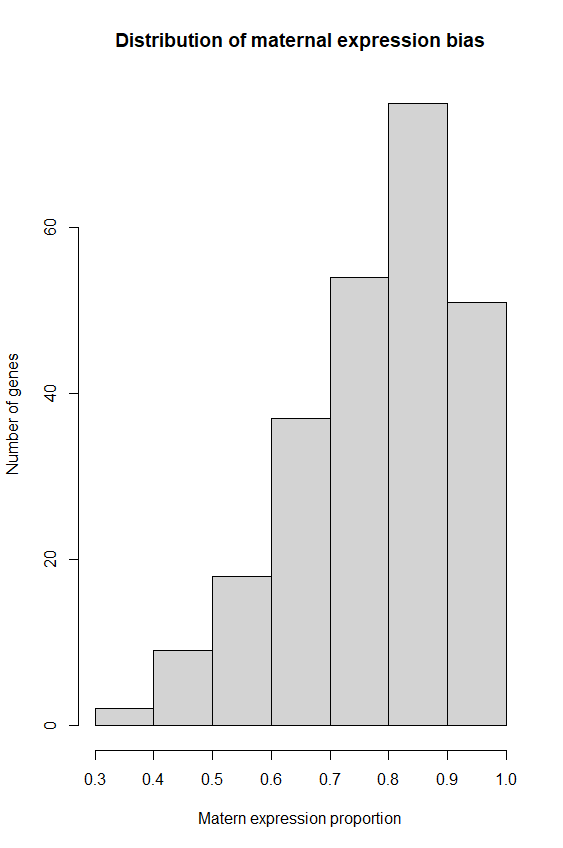
**Figure S3. Distribution of maternal bias of allele expression in males.** Data show both male-biased and unbiased genes for which maternal expression bias is less than 1.0 (completely maternal expression).
